# Interpreting the dynamic pathogenesis of Parkinson’s disease by longitudinal blood transcriptome analysis

**DOI:** 10.1101/2020.10.26.356204

**Authors:** Gang Xue, Gang Wang, Qianqian Shi, Hui Wang, Bo-Min Lv, Min Gao, Xiaohui Niu, Hong-Yu Zhang

## Abstract

Achieving an improved understanding of the temporal sequence of factors involved in Parkinson’s disease (PD) pathogenesis may accelerate drug discovery. In this study, we performed a longitudinal transcriptome analysis to identify associated genes underlying the pathogenesis of PD at three temporal phases. We firstly found that multiple initiator genes, which are related to processes of olfactory transduction and stem cell pluripotency, indicate PD risk to those subjects at the prodromal phase. And many facilitator genes involved in calcium signaling and stem cell pluripotency contribute to PD onset. We next identified 325 aggravator genes whose expression could lead to disease progression through damage to dopaminergic synapses and ferroptosis via an integrative analysis with DNA methylation. Last, we made a systematic comparison of gene expression patterns across PD development and accordingly provided candidate drugs at different phases in an attempt to prevent the neurodegeneration process.

## Introduction

Parkinson’s disease (PD) is a common chronic neurodegenerative disorder characterized by tremor and postural instability in motor symptoms along with non-motor symptoms, including hyposmia, sleeping disorder and cognitive decline^1,2^. The primary pathological features of PD are aggregations of alpha-synuclein (α-syn) in the cytoplasm, as well as the elimination of dopaminergic neurons in the substantia nigra pars compacta (SNc)^1^.

Currently, the pathogenesis of PD has not been thoroughly elucidated, and many factors have been identified to contribute to the neurodegenerative process of PD. Recently, a new conceptual model of PD pathogenesis was proposed, in which these multifaceted factors can be divided into three temporal phases, namely, triggers, facilitators, and aggravators, and they may play distinct roles at different phases of PD^3^. Triggers are agents, such as environmental toxins and viral infections, that initiate the disease process. Accompanying or following triggers, facilitators spread the pathology to the central nervous system, leading to the onset of PD motor symptoms. Facilitators are proposed to include systemic inflammation, mitochondrial dysfunction, and genetic mutations. After PD manifests, aggravators, such as impaired autophagy and neuroinflammation, can directly contribute to disease progression. This conceptual model provides a new definition for PD pathogenesis, but the framework requires further investigation and validation at the molecular level. Elucidating the temporal sequence of pathogenesis in the disease process may be helpful for applying appropriate therapeutic agents when the related factor is most active^3^.

Transcriptome sequencing is a useful technique that is employed to quantitatively investigate genes responsible for the underlying pathogenesis. Previous transcriptomic studies of PD examined gene expression at a single time point^4,5,6^, hindering the observation of expression changes in disease development. By monitoring the same individual at different stages or time points, longitudinal transcriptomics analysis can capture the gene expression patterns during the temporal phases of disease and therefore has the potential to be employed to study the systemic evolution of pathogenesis^7^. The Parkinson’s Progression Markers Initiative (PPMI) is a landmark longitudinal observational study to comprehensively evaluate PD and identify biomarkers of disease progression^8^. In 2019, the PPMI reported the longitudinal RNA resource of whole blood samples, which enabled us to determine the sequence of pathogenesis in PD development.

In this study, we performed a longitudinal transcriptome analysis with blood samples from the PPMI to study the dynamic pathogenesis of PD (Fig. 1). By analyzing the subjects who developed from prodromal to PD, we first identified initiator genes that indicated the risk of disease by survival analysis. Next, facilitator genes were determined to contribute to the onset of PD through differential expression analysis. Moreover, aggravator genes were identified in 252 *de novo* PD patients observed from baseline to the third year. Causal inference and methylome-wide association study (MWAS) provided further validation and insight into aggravation. Based on the distinct gene expression patterns observed in these three phases, we suggested potential therapies to be employed at the appropriate stages to prevent neurodegeneration in PD. Notably, we demonstrated that withaferin A has the ability to promote the differentiation of neural stem cells into dopaminergic neurons *in vitro* experiment. The results suggest great potential of withaferin A to treat prodromal or early stage PD patients.

**Fig. 1.**
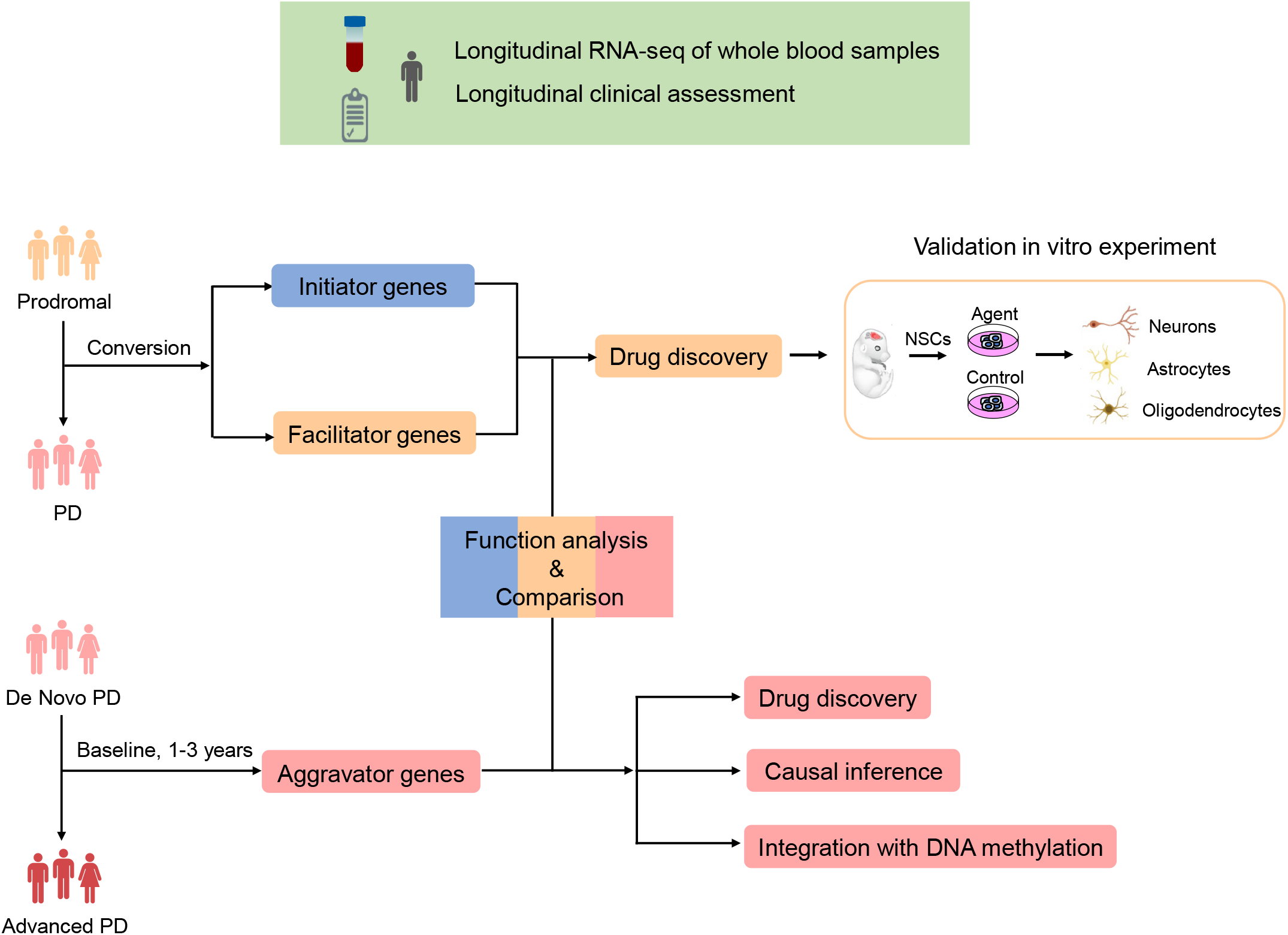
Study workflow. Longitudinal RNA-seq of whole blood samples and corresponding clinical assessment were obtained from Parkinson’s Progression Markers Initiative (PPMI) database (https://www.ppmi-info.org/data). (up) Flowchart of analysis in the conversion process. (right) Flowchart of analysis in the aggravation process.

## Results

### Overview of study

In this study, two cohorts from PPMI were utilized to interpret the dynamic pathogenesis of PD (Fig. 1). Sixty-four prodromal subjects who had a diagnosis of hyposmia or rapid eye movement sleep behavior disorder (RBD) or at least one motor symptom of PD were followed from 2013 to 2019, some of whom had converted to PD during longitudinal observation. The conversion process was employed to infer the pathogenesis of initiation and facilitation. To investigate the pathogenesis of disease aggravation, 252 *de novo*, untreated idiopathic PD patients were selected at baseline and subsequently observed at 1 year, 2 years, and 3 years. In this instance, the Movement Disorder Society-Unified Parkinson’s Disease Rating Scale (MDS-UPDRS) total score was applied to measure the clinical severity of PD.

It is difficult to obtain brain transcriptomes from PD patients, while blood transcriptomes are more accessible and less invasive to obtain from patients, making it easy to dynamically monitor cellular injuries that occur in PD. RNA-seq data were obtained from whole blood samples under the stringent quality controls applied by PPMI (Methods). Notably, ~60-70% of genes expressed in the blood are also expressed in brain tissues, as determined by examining mRNA only, suggesting that the blood may serve as a surrogate tissue for brain transcriptome analysis and facilitate the identification of PD-related molecular signatures.

### Identification of initiators

In the previous conceptual model of PD pathogenesis, triggers are agents or events that initiate disease pathogenesis during prodromal PD. These triggers, such as pathogens and environmental toxins, are exogenous factors that initiate the disease process^3^. However, we attempted to infer endogenous factors at the molecular level by examining initiator genes, which might indicate the risk of disease and are thought to act in the prodromal stage. To identify these genes, we defined prodromal conversion to PD as an “event” and defined time interval of conversion as the “survival time” to perform survival analysis (Methods). Next, genes that were significantly associated with the risk of conversion were considered initiator genes of PD (2,714 genes, log-rank test *P* < 0.05, Supplementary Table 1).

Many initiator genes that influence the risk of PD onset were involved in signaling pathways regulating pluripotency of stem cells, olfactory transduction, and hepatitis C (Fig. 2a). In line with the triggers proposed, hepatitis C, which is an infectious pathogen, is associated with a higher risk of developing PD^9^. In addition, the dysfunction of olfactory transduction is strongly related to hyposmia, which is one of the earliest non-motor symptoms of PD^10^ and is consistent with the clinical symptoms of prodromal subjects, as previously described. Together with the evidence of α-syn aggregates in olfactory^11^, these findings indicated that the olfactory system may be the major initiator site in PD pathogenesis. Notably, *OR8D1*, one of the key genes to help maintain normal olfactory function^12^, ranks first among the initiator genes (log-rank test, *P* = 3.2e-05). Prodromal subjects with lower expression levels of *OR8D1* were at high risk of converting to PD (Fig. 2b). Thus, the expression of *OR8D1* could be employed to help early diagnosis. In PD, ~50% of nigral neurons are lost before the onset of clinical motor symptoms^13^. According to our results, we hypothesize that the lost neurons might not be renewed due to the damaged pluripotency of stem cells, which can influence the neurogenesis process^14,15^.

**Fig. 2.**
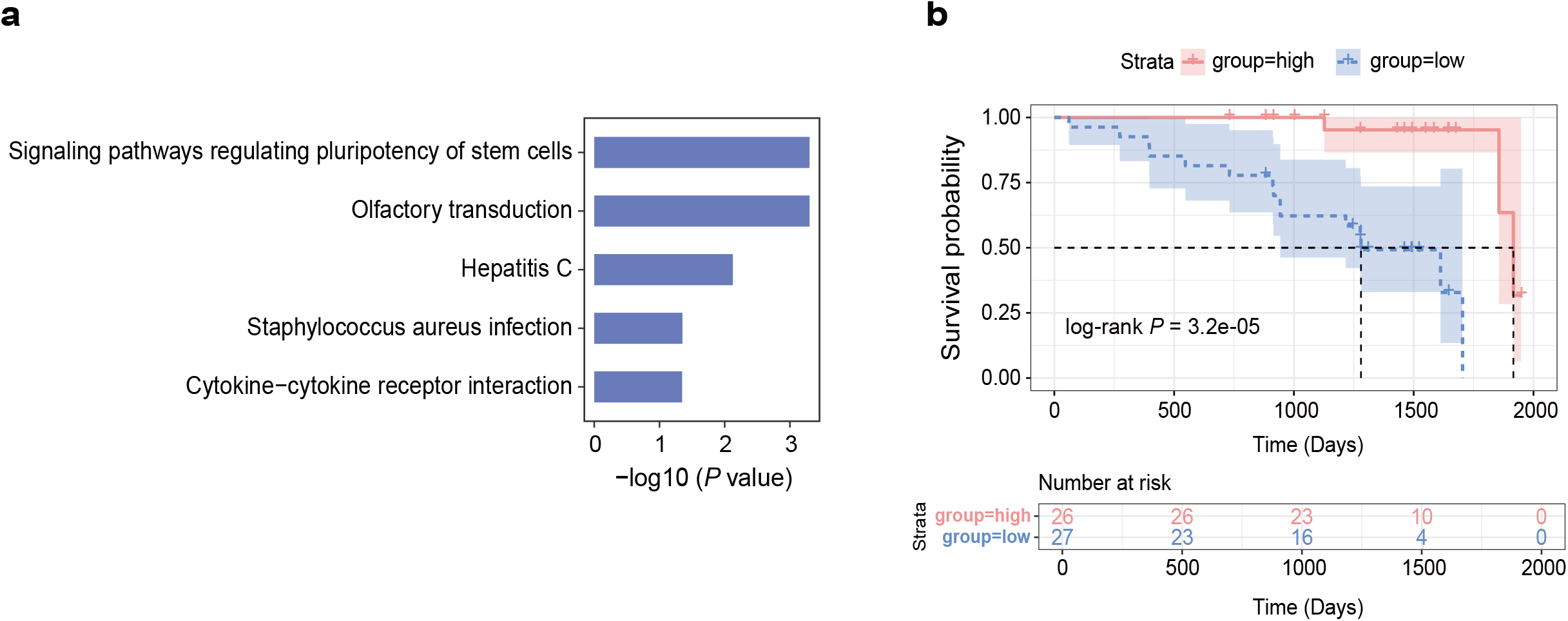
Pathway enrichment and survival analysis of initiator genes. **a** KEGG pathway enrichment of initiator genes by gene set enrichment analysis (GSEA) (genes ranked by −log10(*P*-value)). **b** Kaplan-Meier curves of conversion in prodromal subjects (*N* = 53). The magenta line indicates prodromal subjects with higher expression of *OR8D1*, while the blue line indicates prodromal subjects with lower expression of *OR8D1*, which are at high risk of conversion.

### Identification of facilitators

Facilitators are proposed to help initiators spread PD pathology and damage the central nervous system^3^. Unlike initiator genes, facilitator genes are thought to contribute to the onset of PD and show different expression patterns in prodromal and early-stage PD. To find the facilitator genes, we identified differentially expressed genes (DEGs) by comparing the expression profiles from prodromal to confirmed PD patients in a discovery dataset, which contained 17 RNA-seq samples of five subjects during their natural conversion (Method, Fig. 3a). We identified 1,326 DEGs whose changes in expression during conversion may facilitate the onset of PD (*P* < 0.05, Supplementary Table 2). To assess the robustness of our results, a similar analysis was performed in a validation dataset with 57 prodromal subjects and 390 PD subjects at baseline. Although these two cohorts have no conversion relationship between each other, the large-scale samples can largely eliminate the influence of differences deriving from genetic background in our comparative research. The DEGs in the discovery and validation datasets (false discovery rate (FDR) < 0.01, Supplementary Table 3) had a significant overlap (Fig. 3b), supporting the results in the discovery dataset with limited samples. Therefore, these 1,326 DEGs refer to facilitator genes of PD.

**Fig. 3.**
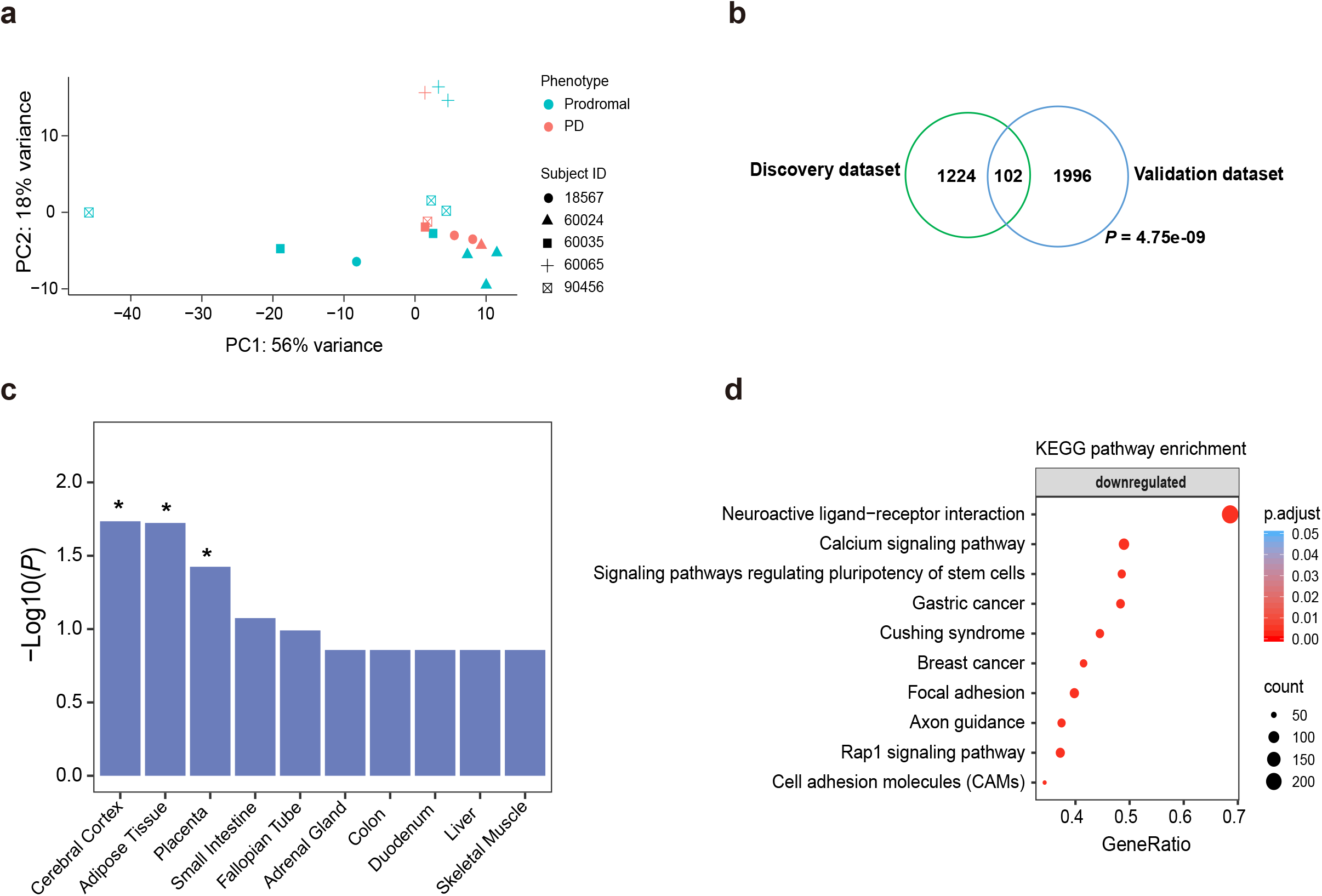
Identifying DEGs between prodromal and PD. **a** PCA clustering plot of RNA-seq data between prodromal and PD. **b** Overlaps of DEGs in the discovery and validation datasets (*P* = 4.75e-09). **c** Tissue enrichment of DEGs between prodromal and PD (\**P* ≤ 0.05). **d** KEGG pathway enrichment of DEGs by GSEA (genes ranked by log2FoldChange).

The facilitator genes were mostly expressed in the cerebral cortex, suggesting that the central nervous system might have been damaged (Fig. 3c). In addition, the facilitator genes were mainly downregulated in the calcium signaling pathway and signaling pathways regulating pluripotency of stem cells during conversion (Fig. 3d). Evidence has shown that changes in calcium (Ca^2+^) levels may trigger a series of events to increase SNc neuronal death sensitivity^16^, including mitochondrial dysfunction, enhanced ROS generation and damaged ATP production. Ca^2+^ homeostasis is also regulated by PD-related proteins, such as α-syn, PINK-1/parkin, leucine-rich repeat kinase 2 (LRRK2), and DJ-1 protein^16^. These findings confirmed the facilitators of a previously conceptual model^3^ and indicated that dysfunction in Ca^2+^ signaling is one of the earliest events in the pathogenesis of PD^16^. We also noted that the pathway of stem cells pluripotency was also captured by initiator genes; thus, the neurogenesis process was influenced in both the prodromal and early stages of PD, which may also imply its causative role in the onset of PD. Taken together, these findings indicated that the dysfunctions in calcium signaling and stem cell pluripotency might be considered facilitators of PD pathogenesis.

### Identification of aggravators

Once PD manifests, aggravators can directly promote the neurodegenerative process and contribute to disease progression. In fact, although there are some interventions for PD, at present, there is no agent that can halt PD progression. Most of the patients showed aggravation within a few years after diagnosis^17^. In addition, consistent with our observation (Supplementary Fig. 1a-b), the clinical manifestations of patients differ greatly in the trajectory of aggravation^18^, hindering efforts to identify a suitable treatment. Although molecular mechanism underlying PD aggravation has critical importance in the long-term prevention and treatment of this disease, it has not been elucidated to date. Therefore, we searched for potential aggravator genes to determine the underlying mechanisms of PD.

With the longitudinal blood transcriptome of 252 patients at four serial time points (baseline, 1 year, 2 years, and 3 years), we constructed linear mixed effects models (Methods) to characterize the association between gene expression change and clinical severity during disease aggravation. Using each subject’s baseline samples as their own control, this analysis reduces the noise derived from the genetic background of patients and improves the precision. We identified 325 genes significantly associated with PD aggravation (FDR ≤ 0.05; Supplementary Table 4), 307 of which were consistently downregulated and 18 of which were upregulated, across the entire aggravation process (Fig. 4a). Taking downregulated aggravator genes as an example, at early stages (baseline and 1 year), these genes were expressed at a relatively high level, but the gene expression level decreased significantly in many patients after 2-3 years. Such an expression trend was also observed in an independent dataset from PD patients (Supplementary Fig. 2), suggesting that the dynamic molecular functions dominating disease aggravation may help the development of drugs to intervene in the process. Moreover, the downregulated aggravator genes were mainly enriched in the dopaminergic synapse pathway, cGMP-PKG signaling pathway, and ferroptosis (Fig. 4b). It is clear that the suppressed function of dopaminergic synapses can damage dopaminergic neurons. Ferroptosis is an iron-dependent autophagic cell death process^19^, and prior research has elucidated its mechanism in PD neurodegeneration. For example, as a key component of ferroptosis, iron generates hydroxyl radicals and induces the oxidation of dopamine, which may lead to an oxidative environment and thus contribute to the death of dopaminergic neurons in the SNc^20^.

**Fig. 4.**
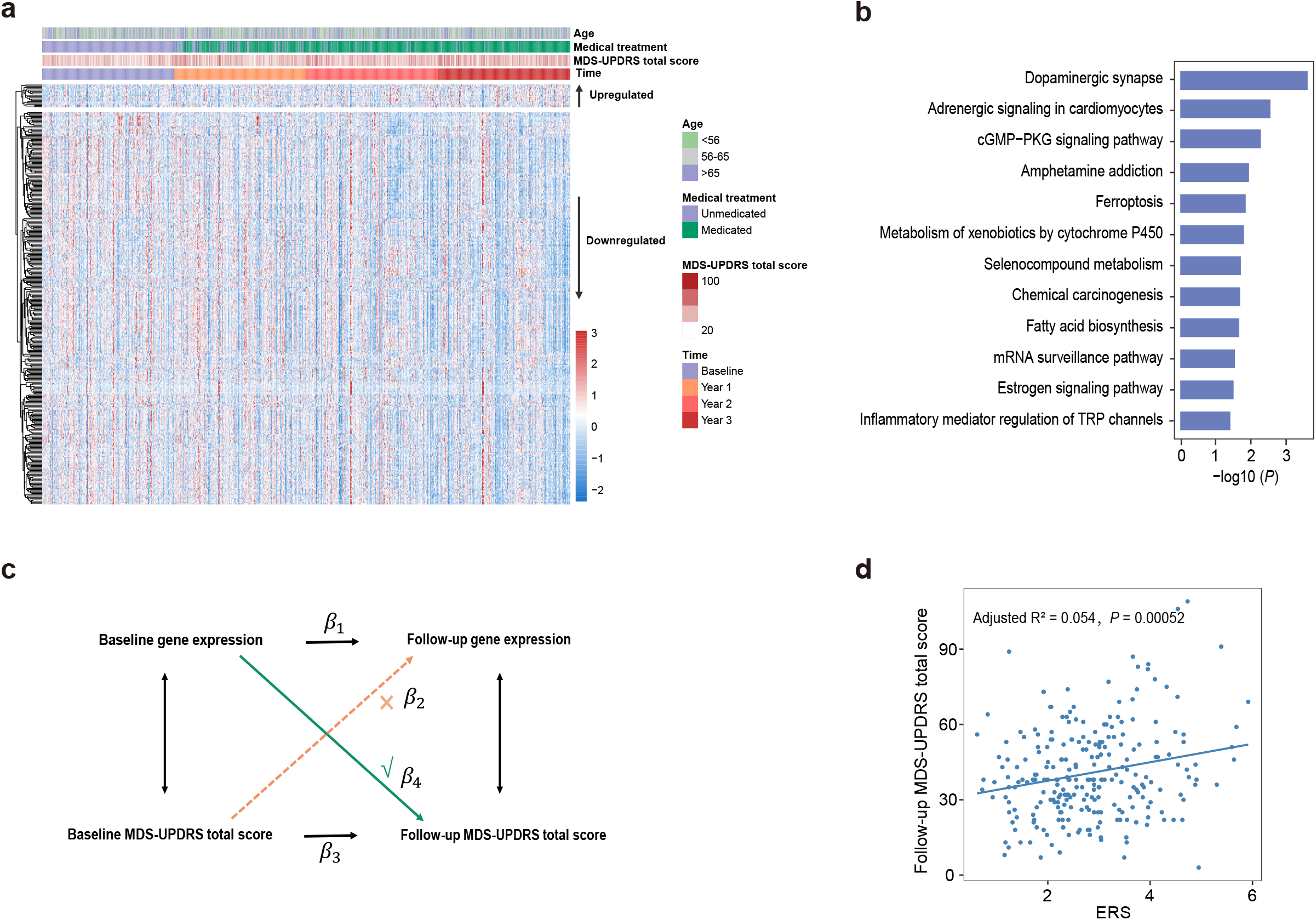
Functional analysis of aggravator genes and causal inference. **a** Heatmap of aggravator genes in longitudinal PD patients (*N* = 252). **b** KEGG pathway enrichment of downregulated aggravator genes. **c** Causal inference based on the cross-lagged path analysis model. *β*_2_ and *β*_4_ are cross-lagged path coefficients. Genes with significant unidirectional paths (*β*_4_, *P* < 0.05) from baseline gene expression to follow-up MDS-UPDRS total score (the third year), while those without significant path coefficients (*β*_2_) from baseline MDS-UPDRS total score to follow-up gene expression are considered to be causing genes in disease aggravation. **d** Correlation between the weighted gene expression risk score (ERS) at baseline and MDS-UPDRS total score at follow-up (the third year).

What remained to be confirmed was whether the changes in the expression levels of aggravator genes were the consequences or causes of PD progression. Therefore, we next performed cross-lagged panel analyses (Methods, Fig. 4c) to investigate the cause-effect relationship between the 325 aggravator genes and clinical severity (i.e., MDS-UPDRS total score). Ten aggravator genes showed significant path coefficients (*β*_4_) from baseline gene expression to follow-up MDS-UPDRS total score (the third year) but did not show significant path coefficients (*β*_2_) from baseline MDS-UPDRS total score to follow-up gene expression, demonstrating that the changes in expression exhibited by these genes may be the causes, rather than the consequences of PD aggravation (Supplementary Table 5). Based on the unidirectional relationship from baseline gene expression to follow-up clinical severity at ten genes, we further examined the association between weighted gene expression risk score (ERS) at baseline and MDS-UPDRS total score at follow-up and found a significant positive correlation between them (Methods), which confirmed that these genes drive the process of aggravation and could be utilized for disease prediction (Fig. 4d).

### Integrative analysis of aggravator genes and DNA methylation

As is well-known, the alterations of gene expression are affected by comprehensive factors, such as genetic variation and epigenetic changes. In this study, we further integrated the baseline blood methylation profiles to elucidate the epigenetic regulation related to disease aggravation. Methylome-wide association study (MWAS) were performed to distinguish the disease signals from CpG methylation (Method). Since methylation alterations may achieve long-term regulation of disease development and do not cause clinical manifestations immediately^21,22^ (Supplementary Fig. 3), we constructed a linear model with the baseline methylome and the rate of PD aggravation. A total of 1,416 CpG sites (*P* ≤ 0.001, Supplementary Table 6) were found to be associated with the rate of aggravation (Fig. 5a), and methylation score derived from these CpG sites accounted for approximately 19% of its variance (Method, Fig. 5b). In addition, previous research identified 138 CpGs that significantly changed over time in PD patients with longitudinal genome-wide methylation data^23^. Although there is no overlap between our aggravator rate-associated CpGs and the previously identified 138 CpGs, the genes mapped by CpGs showed significant overlap (*P* = 7.8e-06, Fig. 5c), further indicating that these 1,416 CpGs identified by us were related to disease aggravation. Moreover, the aberrant methylation events that occurred in many PD-related pathways overlapped with those pathways enriched by aggravator genes, e.g., dopaminergic synapse and ferroptosis, which led us to investigate their functional associations.

**Fig. 5.**
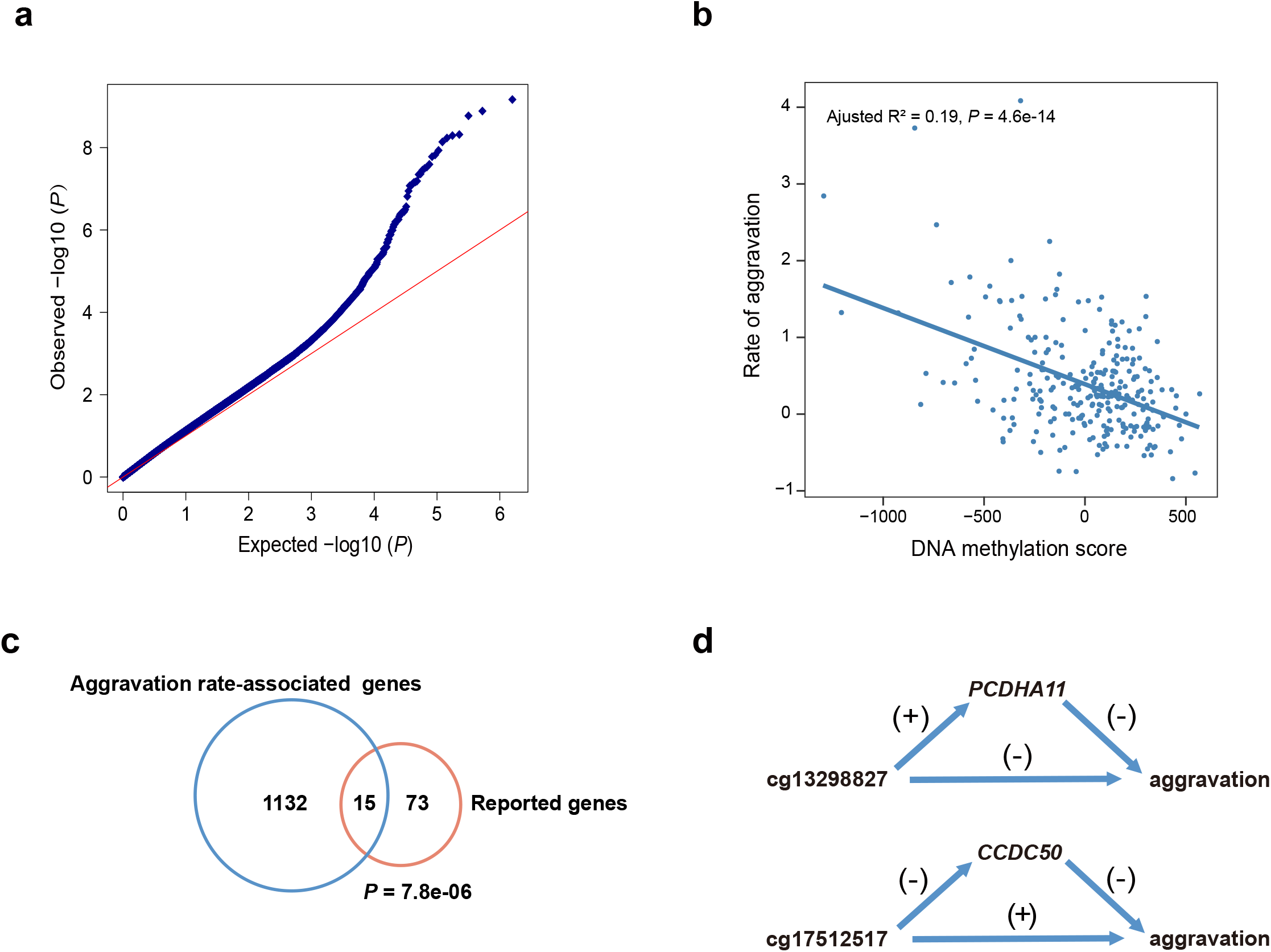
Assessment of the blood methylome and investigation of their regulatory relationship with aggravator genes. **a** Quantile-quantile (Q-Q) plot of MWAS result using rate of aggravation as clinical traits (*N* = 268). **b** Linear relationship between DNA methylation score and rate of aggravation of PD patients. **c** Overlaps of our aggravator rate associated-genes with literature-reported PD aggravator**-**associated genes (CpGs map to genes). **d** (upside) The associations between cg13298827, *PCDHA11* and aggravation; (downside) The associations between cg17512517, *CCDC50* and aggravation. Positive (+) and negative (-) associations are shown in the figure.

We subsequently explored the associations between our aggravator genes and these aggravator rate-associated CpGs (Method) and found that the changes in expression in all aggravator genes exhibited significant relationships with these CpGs (*P* < 0.05). In particular, cg13298827 was positively correlated with expression changes of *PCDHA11* located within +/-10 kb (Fig. 5d). *PCDHA11* may be involved in the creation and maintenance of specific neuronal connections in the brain, and its dysfunction may influence the signaling and communication between neural cells, thereby leading to the aggravation of PD. Since some regulations may have long-range interactions, we next integrated high-throughput chromosome conformation capture (Hi-C) data from the brain dorsolateral prefrontal cortex^24^ (http://resource.psychencode.org) and identified 18 CpG-gene interactions in the topologically associating domains (TADs) (Supplementary Table 7). For example, cg17512517 may influence disease aggravation by negatively regulating expression changes of *CCDC50* (Fig. 5d). Taken together, our results indicated that DNA methylation may regulate the changes in expression levels of aggravator genes, which may modulate PD progression.

### Inferred cellular changes associated with aggravation

In addition to identifying genes associated with aggravation, we also examined which cell types were changed along with PD aggravation. In this study, we applied the xCell algorithm^25^ to infer the abundance scores of 64 immune and stromal cell types from the RNA-seq expression profiles of each PD sample at baseline and after 1-3 years. Analogous to our aggravator gene identification flow, 14 cell types were found to be significantly associated with PD aggravation (FDR < 0.05, Supplementary Fig. 4). Remarkably, neurons, although slightly increased in the third year, significantly decreased overall with disease aggravation, which is in keeping with the loss of neurons observed in the SNc of PD patients^1^. In addition, some of the gene signatures of neurons were also downregulated aggravator genes, such as *NDRG4, PKIA, SHISA6, STX16, ZNF14*, and *ZNF606*, which confirmed our previous results. Nonetheless, it has not been determined whether changes in cellular proportions take place before transcriptomic changes or after them. Moreover, a recent study provided more detailed evidence regarding the cell types underlying the pathogenesis of PD based on single-cell transcriptomic data, demonstrating that cholinergic neurons, monoaminergic neurons, enteric neurons, and oligodendrocytes were related to PD and that these cells changed at the earliest stages of disease progression^26^.

### Distinct expression patterns between initiator, facilitator and aggravator genes

We next investigated how PD develops by tracking the trajectory of gene expression patterns. We first observed that the initiator, facilitator and aggravator genes have little overlap (Fig. 6a), which is reminiscent of the distinct biological processes of these three temporal phases, as previously discussed. When further investigating their connection in the protein-protein interaction (PPI) network, we observed that the facilitator genes possess more degrees than initiator genes and tend to be located in the hub of the network (Fig. 6b-c), while aggravator genes are located in the hub compared to the facilitator genes (Fig. 6d-e). Since the expression changes in initiator genes precede the facilitator genes and the changes in facilitator genes precede the aggravator genes, it appears that the dysfunction first occurred in the signature genes at the early phase, which would further spread the disease disturbance to spur neurodegeneration by giving way to new signature genes, exhibiting dynamic pathogenesis in disease development and implying that although distinct biological functions are presented by the three temporal phase genes, they can connect to one another in various ways. This complex evolution of pathogenesis may account for the difficulties in halting or slowing PD development. Taken together, these results confirmed the difference in pathogenesis among the three temporal phases of PD while also demonstrating the connections among them.

**Fig. 6.**
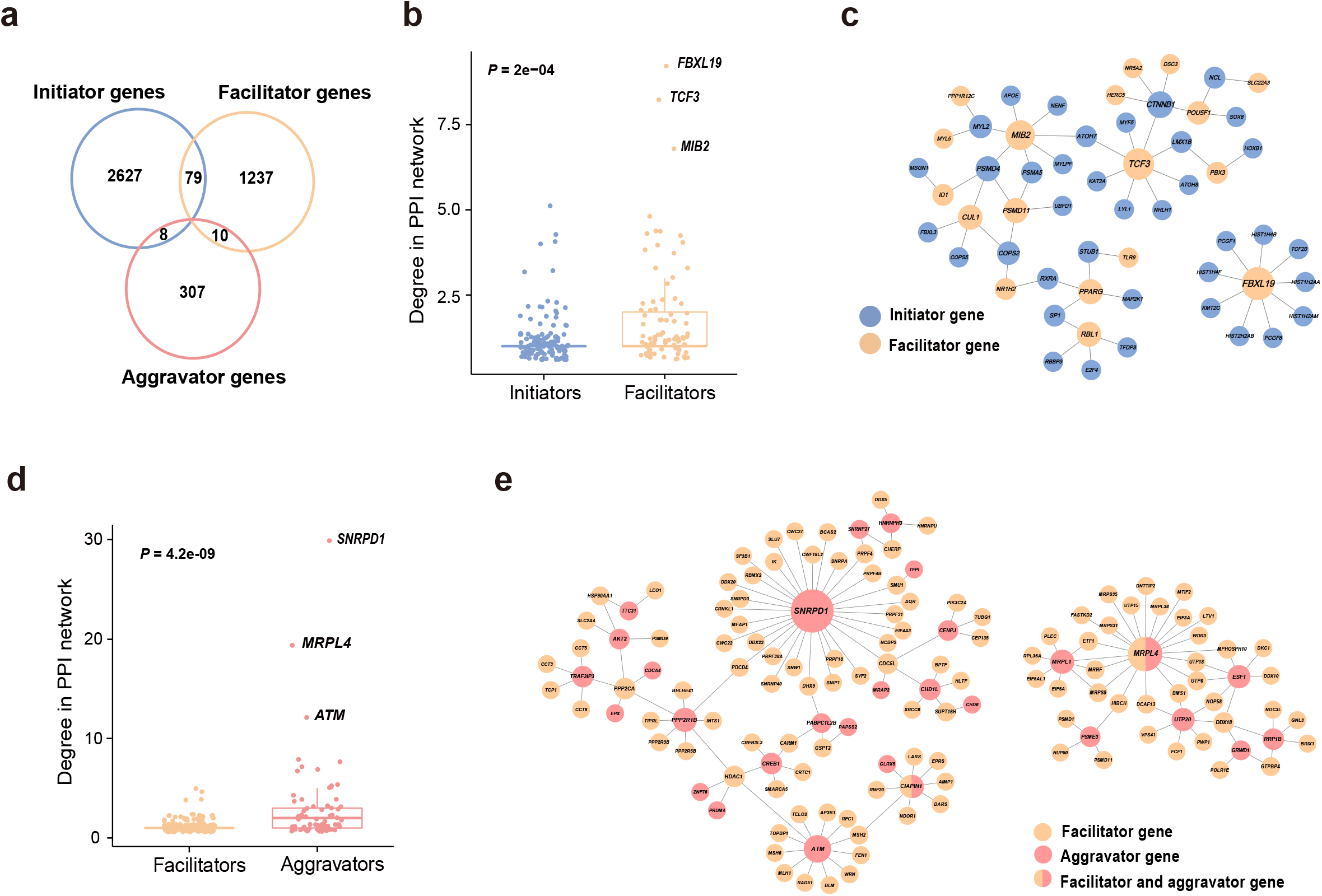
Relationship between initiator, facilitator and aggravator genes. **a** Initiator, facilitator and aggravator genes do not show significant overlaps with each other (*P* > 0.05). **b** Boxplot of the connected degrees of initiator and facilitator genes in the PPI network. Facilitator genes showed more connections than initiator genes (*P* = 2e-04). **c** Network diagram depicting the interaction between initiator and facilitator genes. The facilitator genes (yellow) tend to be located in the hub of the network. **d** Boxplot of the connected degrees of facilitator and aggravator genes in the PPI network. Aggravator genes showed more connections than facilitator genes (*P* = 4.2e-09). **e** Network diagram depicting the interaction between facilitator and aggravator genes. The aggravator genes (pink) tend to be located in the hub of the network.

### Identification of potential drugs for different phases of PD

After elucidating the factors involved in PD pathogenesis in three temporal phases, we next sought to identify potential therapies to reverse or slow the development of PD. Currently, pharmacological treatment and deep brain stimulation are the main therapies to alleviate the symptoms of PD, but they are unable to prevent the neurodegenerative process^27,28^. Another promising therapy is cell-based treatments, which can replace lost neurons with new and healthy neurons to reconstruct disrupted circuits^29,30,31^. We propose that these therapies should be applied at the appropriate phase of PD to maximize the chances of modifying the neurodegeneration process.

As mentioned above, the stem cell pluripotency was influenced in both the prodromal and early stages of PD, and analysis of facilitator genes indicated that the signaling pathways regulating the pluripotency of stem cells (KEGG ID: hsa04550) were downregulated when prodromal subjects were converted to PD patients (Fig. 7a). This result motivated us to seek novel candidate agonists to treat prodromal or early stage PD patients by recovering the function of stem cells and promoting their revitalization of neurons. To this end, L1000CDS^2^ analysis was performed to identify potential agents by upregulating the expression of DEGs in the hsa04550 pathway (Methods, Supplementary Fig. 5a). Next, we obtained three candidate compounds, i.e., atorvastatin, curcumin and withaferin A (WFA), the former two of which have been reported to promote the differentiation of neurons in neural stem cells (NSCs)^32,33^. Moreover, similar results were also obtained in the validation dataset using the same approach (Supplementary Fig. 5a), indicating that WFA is of high confidence for promoting NSCs differentiation into neurons.

**Fig. 7.**
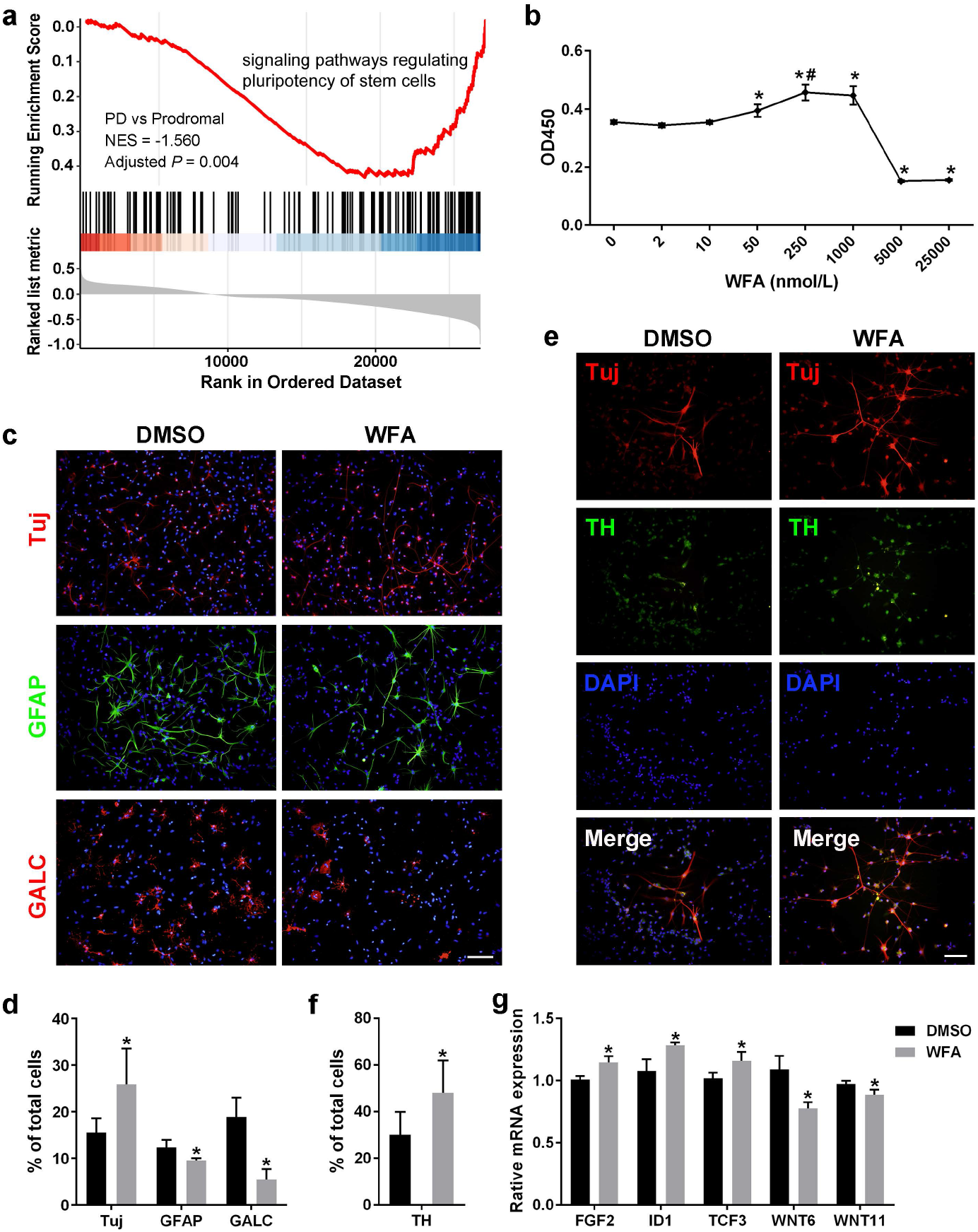
WFA promotes the proliferation and differentiation of NFCs. **a** GSEA plot showing downregulated signaling pathways regulating pluripotency of stem cells when prodromal conversion to PD. **b** Identification of the optimum concentration of WFA on the proliferation of NFCs by the CCK8 test. WFA significantly promoted NSCs proliferation at 50, 250, and 1000 nmol/L (\**P* ≤ 0.05), and the proliferative effects at 250 nmol/L were significantly higher than those at 50 nmol/L (#*P* ≤ 0.05), suggesting that WFA has a dose-dependent effect, whereas no significant difference was observed between 250 nmol/L and 1000 nmol/L, indicating that 250 nmol/L may reach the plateau stage of a dose-dependent effect; thus, 250 nmol/L was adopted to evaluate the effects of WFA on the next NSCs differentiation experiment. **c** Detection of NFCs differentiation by immunofluorescence staining (scale bar =100 μm). **d** Statistical analysis of neurons (Tuj^+^ cells), astrocytes (GFAP^+^ cells) and oligodendrocytes (GalC^+^ cells) between the WFA- and DMSO-treated control groups (\**P* ≤ 0.05). **e** Detection of dopaminergic neurons derived from NSCs using immunofluorescence staining (Scale bar = 100 μm). **f** Statistical analysis of the proportion of dopaminergic neurons (TH^+^ Tuj^+^ cells) in neurons (Tuj^+^ cells) between the WFA and control groups (\**P* ≤ 0.05). **g** Assessment of the mRNA expression of genes regulating the pluripotency of stem cells by quantitative RT-PCR (\**P* ≤ 0.05).

To verify our inferences, we first evaluated the effects of WFA on the proliferation and differentiation of NSCs *in vitro* (Methods). The optimum concentration for NSCs differentiation was identified by the CCK8 test (Fig. 7b). Since NSCs will start to differentiate spontaneously in the absence of basic fibroblast growth factor (bFGF)^34^, we applied WFA or DMSO (control) to treat NSCs without bFGF, and we assessed differentiation by immunofluorescence staining. We observed that WFA can promote NSCs to differentiate more neurons (Tuj^+^ cells), fewer astrocytes (GFAP^+^ cells) and oligodendrocytes (GalC ^+^ cells) compared with the DMSO-treated control group (Fig. 7c-d), which demonstrated that WFA can promote NSCs to differentiate into neurons.

Given that the loss of dopaminergic neurons is an important pathological change in PD^1^, we performed further experiments and observed that the proportion of dopaminergic neurons (TH^+^ Tuj^+^ cells) in neurons (Tuj^+^ cells) under WFA treatment was significantly higher than that in the control (Fig. 7e-f), suggesting that WFA can effectively promote NSCs to differentiate into dopaminergic neurons. Moreover, five WFA-regulated genes were identified from the DEGs (Supplementary Fig. 5a) and further validated by quantitative RT-PCR (Supplementary Table 8, Fig. 7g). Specifically, *FGF2, ID1*, and *TCF3* were upregulated, which is consistent with our hypothesis that NSCs differentiation is promoted by upregulating the DEGs. While *WNT6* and *WNT11* were downregulated, we speculated that the difference between blood tissue and NSCs may lead to this inconsistency. Notably, *TCF3* is a transcriptional regulator involved in the maintenance of neural stem cells^35^; thus, its functional recovery probably promotes the revitalization of neurons. *TCF3* is also a hub node in the network of initiator-facilitator genes (Fig. 6c). Taken together, these results strongly indicated that WFA has considerable potential to cure prodromal and early-stage PD by promoting the differentiation of NSCs into dopaminergic neurons.

With the aggravation of PD, we found that biological terms, such as dopaminergic synapse and the cGMP-PKG signaling pathway, were also enriched by the drug targets of PD, implying their connections in a biological context. Among our aggravator genes, six downregulated aggravator genes were also drug targets of PD, and three of them have agonists in the DrugBank database, which might be considered viable drugs in attempts to slow PD aggravation (Table 1). For example, by upregulating *DRD3* in the dopaminergic synapse pathway, levodopa may be suitable to treat patients with relatively more severe symptoms. In fact, levodopa is effective in controlling most motor symptoms and remains the gold standard in the drug treatment of PD^36^. Moreover, to yield more effective disease-modifying treatments, some of these agonists should be combined to target the various pathogeneses in aggravation. In addition, although *SLC18A1* is not a target of PD to date, it is located in the dopaminergic synapse pathway and connected to known drug targets by dopamine transduction. The downregulation of *SLC18A1* expression may lead to dysfunction of the synaptic vesicle circle, thereby accelerating the severity of the disease (Supplementary Fig. 5b); therefore it may be reasonable to consider *SLC18A1* as a potential target for the design of therapeutic interventions in the future.

**Table 1.** Downregulated aggravator genes and the involved agonists.

| Aggravator genes | Agonists (DrugBank) |
| --- | --- |
| *ADRA2C* | Brimonidine, Norepinephrine, Droxidopa,<br>Clonidine, Bromocriptine, Apomorphine,<br>Paliperidone, Ephedra sinica root, Metamfetamine,<br>Xylometazoline, Agmatine, Oxymetazoline, Racepinephrine |
| *DRD3* | Ropinirole, Pramipexole, Apomorphine,<br>Cariprazine, Pergolide, Bromocriptine,<br>Lisuride, Cabergoline, Rotigotine,<br>Dopamine, Levodopa, Captodiamine,<br>Dihydro-α-ergocryptine, Tianeptine |
| *OPRD1* | Codeine, Dextropropoxyphene, Hydrocodone, Morphine,<br>Butorphanol, Fentanyl, Etorphine, 3-Methylthiofentanyl,<br>Dimethylthiambutene, Loperamide, 3-Methylfentanyl,<br>Oxycodone, Diamorphine, Tramadol, Methadone, Sufentanil,<br>Carfentanil, Amitriptyline, Diphenoxylate, Levorphanol,<br>Dihydromorphine, Remifentanyl, Ketobemidone,<br>Dextromethorphan, Opium, Loxicodegol, Butyrfentanyl,<br>Met enkefalin, Hydromorphone |

## Discussion

In this study, we identified distinct molecular patterns in the development of PD with a longitudinal blood transcriptomic analysis and provided a modified conceptual model for PD pathogenesis. The results of this study helped us to identify appropriate therapies for PD patients at different stages of the disease.

At the prodromal stage of PD, we found that olfactory-associated genes were first reflected in the initiation of the disease and subsequently provided a simple and effective method to help early diagnosis by examining the expression of the olfactory receptor *OR8D1* in blood samples. Since hyposmia is also a clinical feature of early-stage patients^10^, our results could be coupled with clinical observations. Nonetheless, the validity of these results should be confirmed in larger prospective experiments. In addition, the expression of the initiator genes also supported that hepatitis C is a trigger to spark the disease^3,9^.

After the prodromal subjects were diagnosed as PD patients, the expression of the facilitator genes indicated that dysfunction of calcium regulation might contribute to neuronal death, and the neurons were unable to be renewed due to the decreased pluripotency of stem cells. Changes in calcium levels can induce a series of events, including mitochondrial dysfunction, enhanced ROS generation, and damaged ATP production^16^. Calcium is also regulated by PD-related proteins, such as α-syn and LRRK2^16^. These findings confirmed the facilitators of the previously proposed conceptual model^3^.

With the progression of PD, the expression of the aggravator genes showed that the dysfunction of dopaminergic synapses, cGMP-PKG signaling, and ferroptosis may represent the main factors of the neurodegeneration process. Ferroptosis is an autophagic cell death process that contributes to autophagy activation^19^. In line with the role played by aggravators in a previous conceptual model, these findings highlight the importance of autophagy in disease aggravation^3^. Other aggravators identified by this study need to be confirmed through further experiments. In addition, we noted that although the expression levels of aggravator genes are significantly changed with progression, the magnitude of these expression level changes was not dramatic (|log2 (third year/baseline)| ≤ 1). One of the reasons for this finding may be that PD injury occurs in brain tissue; thus, signal intensity may be reduced when it reaches the bloodstream. The blood environment may also possess biological signals from the internal environment or other organs, which may interfere with the extraction of disease signals. In addition, PD is a chronic disease; therefore, gene expression does not change dramatically, which also highlights the necessity of conducting longitudinal studies that can capture this moderate dynamic change. Interestingly, by integrating DNA methylation, we observed that methylation alterations at baseline did not cause clinical manifestations immediately but may have influenced the PD aggravation rate by regulating the changes in expression levels of aggravator genes. A similar phenomenon was also reported by other chronic conditions^21^, making it possible to detect and monitor PD aggravation earlier. However, it has not been determined how these CpGs regulate gene expression changes and how long they are sustained.

According to the pathogenesis involved at different stages of PD, we further inferred and demonstrated that WFA has the ability to promote the differentiation of neural stem cells into dopaminergic neurons with strong potential to treat prodromal or early stage PD patients (Fig. 7b-g). WFA is a major active component of *Withania somnifera*, which has been demonstrated to protect dopaminergic neurons in Parkinsonian mouse models and has exhibited a strong antioxidative effect^37^. Moreover, *Withania somnifera* was also reported to treat other disorders related to the central nervous system^37^; thus, the future research should investigate whether WFA has a similar mode of action in the treatment of other diseases, such as Alzheimer’s disease. For advanced PD patients, we proposed a list of drugs, such as levodopa, droxidopa, and dextromethorphan, that may be employed to slow PD aggravation. The appropriate drug combination addressing multiple targets and pathogenesis would benefit severe patients. Moreover, our results may also lead to the reconsideration of previously failed clinical trials, which may have been performed in nonspecific PD patients^3^.

## Methods

### Clinical data analysis of PD patients

In this study, the Movement Disorder Society-Unified Parkinson’s Disease Rating Scale (MDS-UPDRS) total score, which relates to various aspects, including non-motor and motor symptoms, as well as motor complications^38^, was applied to measure the severity of PD patients. A higher score indicates an individual with a more serious condition. For the use of PD medications, there are seven medical treatments defined in PPMI, including unmedicated, levodopa, and levodopa plus dopamine agonist. In this study, we simply classified patients as unmedicated and medicated for subsequent analyses; for age characteristics, patients were classified into three groups adopted from PPMI: group 1 (< 56 years), group 2 (56-65 years), group 3 (> 65 years). Missing values of demographic and clinical information were imputed using the median value.

### Omics data and quality control

For RNA-seq data of whole blood samples, quality control and quantification of gene or transcript expression were applied by PPMI. Briefly, FASTQ files were merged and aligned to GRCh37 (hs37d5) by STAR (v2.4K)^39^. The read counts of genes or transcripts were calculated by FeatureCounts^40^ and abundance estimates (transcripts per million, TPM) via salmon^41^. Samples with obvious quality control issues, such as gender mismatch, principal component analysis (PCA) outliers and low read counts, were filtered. After quality control, most samples have a mean Spearman correlation (> 0.87) to all other samples. To reduce the influence of technical noise, we removed genes with counts/TPM of 0 in more than 80% of the samples in the subsequent analysis. Importantly, according to PPMI analysis, ~60-70% of genes in whole blood are also expressed in brain tissue by examining mRNA expression only. Detailed information regarding the RNA sequencing and quality control can be found on the PPMI website (http://ppmi-info.org/).

For DNA methylation (DNAm) data of whole blood samples, PPMI applied the Illumina Human Methylation EPIC array (alias 850K methylation array) to assess methylation status over 850,000 methylation sites across the whole genome. The β values for each probe, which ranged from 0 (no methylation) to 1 (100% methylation), were utilized to measure the methylation level of the targeted CpG site. Quality control included removing low-quality samples and probes, as well as normalizing blood cell composition. On that basis, we also applied the following filtering criteria in this study^42^: removal of probes targeting X and Y chromosomes (*N* = 19,701), removal of cross-reactive probes on the EPIC array (*N* = 43,254), and removal of probes overlapping genetic variants at targeted CpG sites (*N* = 12,510), leaving 792,150 CpG sites for the subsequent analysis. CpGs mapped to genes according to GPL21145 from Gene Expression Omnibus (GEO, http://www.ncbi.nlm.nih.gov/geo/).

### Identifying initiator genes

Sixty-four prodromal subjects were recruited at baseline with a longitudinal observation beginning in 2013, and 18 prodromal subjects converted to PD by 2019. After matching RNA-seq samples and removing subjects without longitudinal observation, 53 prodromal subjects were kept for the downstream analysis. To identify initiator genes of PD, we defined prodromal conversion to PD as an “ event” and defined the time interval from prodromal to PD as the “survival time” to perform survival analysis. For each gene, baseline prodromal subjects were divided into two groups based on the median expression of the gene. Next, survival curves were constructed by Kaplan-Meier survival curves, and the log-rank test was applied to compare whether there was a significant difference in the overall conversion between the two groups using the R package survival.

### Identifying facilitator genes

To identify facilitator genes, we first performed differential expression analysis in a discovery dataset, which contained 17 RNA-seq samples of five subjects after matching clinical status and RNA-seq samples. Specifically, the heterogeneity between subjects is an important confounding factor identified by PCA (Fig. 3a); thus, we applied a multi-factor design that included the subjects and their condition (prodromal and PD) information in the design formula to identify DEGs using the R package/Bioconductor DESeq2^43^. This design will account for differences between the subjects when estimating the effect due to the condition. The validation dataset contained 57 prodromal and 390 PD RNA-seq samples at baseline without a conversion relationship between each other; therefore, the DEGs of the validation dataset were identified only by estimating the effect due to the condition.

### Identifying aggravator genes

In this study, 252 PD patients both included clinical and RNA-seq data at four serial time points (baseline, 1-3 years). We first filtered the genes whose expression did not significantly change during the four time points based on repeated measures ANOVA with the R package ez. The *P*-value was adjusted by sphericity corrections followed by multiple testing using FDR. Genes with FDR ≤ 0.01 were kept for subsequent analyses (3435 genes). The linear mixed-effects model was constructed to identify which gene was associated with PD aggravation using the *lme* function in the R package nlme. Each model included the MDS-UPDRS total score as the dependent variable; gene expression, age, and time (i.e., baseline, 1-3 years), and medication (unmedicated or medicated) as fixed effects; Each subject variation was included as a random effect and considered their different response to medical treatment, therefore each model was constructed as follows:

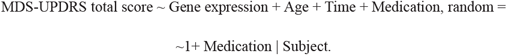

Genes with FDR ≤ 0.05 were defined as aggravator genes (325 genes, Supplementary Table 4). The *β* coefficient of the gene in the model lower than 0 indicates that it was downregulated with PD aggravation; otherwise, it was upregulated. The R package Pheatmap was subsequently used to plot the heatmap of aggravator genes. Notably, some PD samples were abandoned in the above analysis because they did not possess RNA-seq data for four serial years. In fact, most of them were measured three times, involving 425 RNA-seq samples, which were regarded as an independent dataset to assess the robustness of the aggravator genes.

### Cross-lagged panel analysis

Cross-lagged panel analysis was used to investigate cause-effect relationship between the expression of aggravator genes and clinical severity (Fig. 4c). Briefly, baseline gene expression and MDS-UPDRS total score were adjusted for age by regression residual analyses and then standardized with Z-transformation (mean = 0, SD = 1). The same process was performed in the follow-up year (third year) with additional adjustment for the times of medical treatment. Then, two following equations were built:

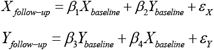

Here, *X* is aggravator gene and *Y* is MDS-UPDRS total score. The linear regression coeffcients *β*_1_ and *β*_3_ describe the autoregressive effects, and the regression coeffcients *β*_2_ and *β*_4_ represent cross-lagged effects. All parameters and statistics were estimated by the *sem* function in the R package lavaan based on these two equations. Next, the weighted gene expression risk score (ERS) was calculated as follows:

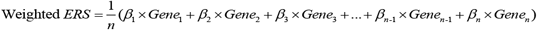

where *β* is the cross-lagged path coefficient (*β*_4_) for each gene, and _*n*_ is the number of genes included.

### MWAS analysis

A linear regression model was constructed to assess the associations between each CpG’s methylation level (β value) and clinical traits of PD at baseline (*N* = 330). The model used for each CpG was as follows:

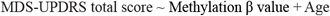

Since variation of DNA methylation (DNAm) may arise prior to disease^21,22^, and we hypothesized that DNAm may influence the aggravation of PD in a long-term manner, we applied the rate of PD aggravation, i.e., the change of MDS-UPDRS total score from baseline to third years, as clinical traits of PD to identify the aggravation rate-associated CpGs. After matching longitudinal clinical and DNAm data, 268 PD patients were retained for the subsequent analysis. Using the *lm* function in R, the linear regression model used for each CpG was as follows:

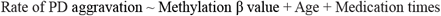

The quantile-quantile (Q-Q) plot was used to diagnose the above MWAS results using the R package qqman. To estimate how much of the rate of PD aggravation could be explained by blood DNAm, the top association CpGs were selected first (*P* ≤ 0.001, Supplementary Table 6). Next, PCA was performed on these CpGs, and each patient was scored by the first principal component using the R package psych. A linear model was constructed to measure the relationship between the score calculated by these CpGs and the rate of PD aggravation.

### Functional analysis

KEGG pathway enrichment and gene set enrichment analysis (GSEA) analysis (2000 total permutations) of gene sets were performed by the R/Bioconductor package clusterProfiler^44^. Tissue enrichment analysis of DEGs between prodromal and PD was investigated by R package/Bioconductor TissueEnrich^45^.

### Integrating transcriptome and methylome on PD aggravation

The RNA-seq profiles and DNA methylation array data were pairwise integrated (*N* = 191) to study the regulatory relationship between each aggravator gene and each aggravation rate-associated CpG site. Each linear regression model included delta gene expression level change from baseline to third years as the dependent variable (log2 (third year/baseline)), β value of a CpG site at baseline as the independent variable, and age and medication times as covariates:

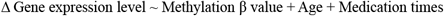

### xCell analysis

xCell^25^ was used to estimate the abundance scores of 64 immune and stromal cell types from the RNA-seq profiles of each PD patent over time (i.e., baseline, 1, 2, 3 years), the abundance scores was then standardized with Z-transformation (mean = 0, SD = 1) Similar to our aggravator genes analysis, linear mixed-effects model was used to inferred which cell type was associated with PD aggravation:

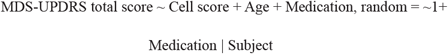

### Observation of initiator, facilitator and aggravator genes in the PPI network

The interactions between initiator, facilitator and aggravator genes were examined in a protein-protein interaction (PPI) network. The PPI network was derived from the STRING database (http://www.string-db.org), and the interaction with experimental score higher than 400 was considered reliable. Notably, since it difficult to observe the network relationship between aggravator genes and facilitator genes identified from the discovery dataset, we merged the facilitator genes both from the discovery and validation datasets to observe their connections with aggravator genes in the PPI network. The gene networks were subsequently visualized using Cytoscape^46^.

### Drug discovery by L1000CDS^2^ analysis

The Library of Integrated Network-based Cellular Signatures (LINCS) program has generated more than 1 million transcriptional profiles to elaborate the human cell expression change under different perturbations of agents^47^. The L1000 dataset is part of LINCS, which is based on the expression of only ~1000 genes^48^. L1000CDS2 is a LINCS L1000 characteristic direction signature search engine that links drugs, genes, and diseases by comparing the input signatures to the background (http://amp.pharm.mssm.edu/L1000CDS2/#/index). In this study, we applied L1000CDS^2^ to predict potential small molecular compounds that may reverse the expression of DEGs between prodromal and PD subjects.

In the discovery dataset, 50 agents were returned by L1000CDS2 according to eight DEGs (*P* < 0.05) in pathways of regulating pluripotency of stem cells (KEGG ID: hsa04550). We further selected three candidate compounds, i.e., atorvastatin, curcumin and withaferin A (WFA), which were used to treat patients with mental disorders in clinical trials (https://clinicaltrials.gov/), indicating the ability of delivery across the blood brain barrier. Using the same approach, WFA was also identified in the validation dataset (57 prodromal and 390 PD RNA-seq samples). To identify potential genes regulated by WFA, we further identified six DEGs in the discovery dataset that overlapped with background genes of WFA in the L1000 dataset, and five of these six potential WFA-regulated genes in the discovery dataset were confirmed by the validation dataset (Supplementary Fig. 5a).

### Isolation and culture of rat fetal neural stem cells

Rat fetal NSCs were isolated and cultured from brain tissues of Sprague-Dawley rats at 14-18 days post-coitum. Briefly, brain tissues were cut into small blocks, digested by pre-warmed StemPro^®^ Accutase (A11105-01, Gibco, NY, USA), filtered by a 70-μm cell strainer (352350, BD Biosciences, NY, USA) and centrifuged. The cell pellet was resuspended in pre-warmed StemPro^®^ NSC SFM complete medium (consisting of KnockOut D-MEM/F-12 with 2% StemPro^®^ Neural Supplement, 20 ng/mL bFGF, 20 ng/mL EGF, and 1% GlutaMAX-I; A1050901, Gibco, NY, USA), inoculated in a dish at 1×10^7^/mL and incubated at 37°C, 5% CO2 and 90% humidity. The NSCs that developed into neurospheres were passaged after 5-7 days (Supplementary Fig. 6a). In subculture, the NSCs were digested by pre-warmed StemPro^®^ Accutase and plated in dishes precoated with 20 μg/mL fibronectin (33016015, Gibco, NY, USA) to adherent culture at 37°C, 5% CO2 and 90% humidity (Supplementary Fig. 6b). The NSCs were passaged again when 75-90% confluent. The cells were identified by immunofluorescence staining of Nestin at passage 2 (P2) ( Supplementary Fig. 6c-d). All WFA effects research experiments were carried out on cells at passages 2 to 3 (P2~3) to minimize the experimental deviations.

### Analysis of WFA on NSCs proliferation by CCK8 assay

WFA (HY-N2065, MCE, NJ, USA) was dissolved in DMSO to make a 1 mmol/L stock solution, and subsequently, we made culture media containing different concentrations of WFA with StemPro^®^ NSC SFM complete medium. The working concentrations of WFA were 0, 2, 10, 50, 250, 1000, 5000, and 25000 nmol/L. After that step, rat fetal NSCs at P2 were collected and resuspended in culture media containing WFA at 5×10^4^/mL and then seeded at 100 μL/well in 96-well plates immediately. To minimize the experimental deviations, culture medium containing WFA but without NSCs was used as a blank control for each concentration. After incubating the cells at 37°C, 5% CO2 and 90% humidity for 48 hours, 10 μL CCK8 reagent (HY-K0301, MCE, NJ, USA) was added to each well and incubated for 4 hours. The absorbance was measured at 450 nm using a microplate reader (Multiskan MK3, Thermo Fisher Scientific, MA, USA). The working concentration of WFA with the highest average absorbance was used to analyze the effect of WFA on NSCs differentiation.

### Intervention of NSCs differentiation by WFA

Rat fetal NSCs at P3 were plated as an adherent culture on chamber slides (177380, Thermo Fisher Scientific, MA, USA) or 6-well plates pre-coated with 20 μg/mL fibronectin for 1 hour. To investigate the effect of WFA on NSCs differentiation, we withdrew bFGF from StemPro^®^ NSC SFM complete medium and added WFA stock solution (1 mmol/L in DMSO, 1:4000; according to the experimental results of CCK8 assay, the work concentration is 250 nmol/L) or DMSO (D2438, 1:4000, Sigma, MO, USA) into the media. Then, the cells were incubated at 37°C, 5% CO2 and 90% humidity until subsequent immunofluorescence and quantitative RT-PCR detection.

### Analysis of WFA on NSCs differentiation by immunofluorescence staining

After intervention by WFA or DMSO for 48 hours, NSCs undergoing differentiation were fixed, permeabilized and blocked and then incubated with primary mouse anti-Tuj antibody (ab78078, 1:1000, Abcam, Cambridge, UK), primary mouse anti-GALC antibody (MAB342, 1:100, Millipore, MA, USA) or primary rabbit anti-GFAP antibody (16825-1-AP, 1:2000, ProteinTech, Wuhan, China) overnight at 4°C. After washing with PBS, cells were incubated with Alexa Fluor 594 conjugated goat anti-mouse IgG (H+L) cross-adsorbed secondary antibody (A-11005, 1:200, Thermo Fisher Scientific, MA, USA) or Alexa Fluor 488 conjugated goat anti-rabbit IgG (H+L) cross-adsorbed secondary antibody (A-11008, 1:200, Thermo Fisher Scientific, MA, USA) at 37°C for 30 min. Finally, cells were washed with PBS, dyed with DAPI (Roche, Basel, Switzerland), mounted with 50% glycerol/PBS and photographed under a fluorescence microscope (BX51, Olympus, Tokyo, Japan).

After treating NSCs with WFA or DMSO for seven days, double immunofluorescence staining of Tuj and tyrosine hydroxylase (TH) was used to analyze whether WFA could further affect NSCs differentiation into dopaminergic neurons. Briefly, after fixation, permeabilization and blocking, cells were incubated with both primary mouse anti-Tuj antibody and primary rabbit anti-TH antibody (ab112, 1:1000, Abcam, Cambridge, UK) overnight at 4°C. Then, the cells were incubated with mixed secondary antibodies of Alexa Fluor 594-conjugated goat anti-mouse IgG and Alexa Fluor 488-conjugated goat anti-rabbit IgG for 30 min. Finally, the cells were dyed with DAPI, mounted with 50% glycerol/PBS and photographed under a BX51 fluorescence microscope.

### Quantitative RT-PCR

Total RNA was extracted from NSCs treated with WFA or DMSO for 48 hours with TRIzol reagent (15596026, Thermo Fisher Scientific, MA, USA) and reverse transcribed into cDNA using the RevertAid RT Reverse Transcription Kit (K1691, Thermo Fisher Scientific, MA, USA). Then, qRT-PCR mixtures were prepared with the KAPA SYBR^®^ FAST Universal qPCR Kit (KK4601, Sigma, MO, USA) according to the manufacturer’s instructions and run in LightCycler 96 (Roche, Switzerland). The mRNA expression of genes regulating the pluripotency of stem cells was normalized to the expression of GAPDH. The results were compared by the 2-ΔΔCt method. The specific primers (Supplementary Table 8) were synthesized by Tsingke Biological Technology (Wuhan, China).

### Statistical analysis of NSCs experiments

Image-Pro Plus 6.0 software (Media Cybernetics, MA, USA) was used for analysis and cell counting of the fluorescence images. The results from different groups were compared using Student’s *t*-test or one-way ANOVA followed by Tukey’s multiple comparison tests. *P*-value ≤ 0.05 was considered to be significant. All statistical analyses were performed with GraphPad Prism 6 software (GraphPad Software, CA, USA).

### PD drugs and targets collection

The drugs for PD were obtained from clinical trials (release 2019.4.22, at least phase 2 clinical trial drugs, https://clinicaltrials.gov/), and the drug targets were extracted from DrugBank^49^, DGIdb^50^ and TTD^51^. We finally collected 104 PD drugs corresponding to 363 target genes (Supplementary Table 9).

## Data availability

Data used in this study were obtained from the Parkinson’s Progression Markers Initiative (PPMI) database (www.ppmiinfo.org/data). For up-to-date information on the study, please visit www.ppmiinfo.org.

## Acknowledgments

We would like to acknowledge PPMI - a public-private partnership, which is funded by the Michael J. Fox Foundation for Parkinson’s Research and funding partners, including Abbvie, Allergan, Amathus Therapeutics, Avid Radiopharmaceuticals, Biogen, BioLegend, Bristol-Myers Squibb, Calico, Celgene, Denali, GE Healthcare, Genetech, GlaxoSmithKline, Golub Capital, Handl Therapeutics, Insitro, Janssen Neuroscience, Lilly, Lundbeck, Merck, Meso Scale Discovery, Pfizer, Piramal, Prevail, Roche, Sanofi Genzyme, Servier, Takeda, Teva, UCB, Verily, and Voyager Therapeutics. We also thank the National Natural Science Foundation of China (grant 31670779) for supporting this study.

## Author contributions

H.Y.Z. and X.H.N. conceived the project; G.X. collected and performed data analysis with help from X.H.N., Q.Q.S., M.G., and H.W; G.W. performed the WFA experiment. G.X., G.W., Q.Q.S., H.Y.Z., X.H.N., H.W., and B.L wrote and edited the manuscript.

## Competing interests

The authors declare no competing interests.

## Supplementary Figure

**Supplementary Fig. 1 Clinical aggravation of PD patients. a** The overall aggravation of PD patients (*N* = 252). Although most patients are under drug treatment after enrollment, the disease condition is still in progression (Repeated measures ANOVA *P* = 9.15e-20). **b** The aggravation trajectory for each PD patient.

**Supplementary Fig. 2 Heatmap of aggravator genes in independent PD patients (425 RNA-seq samples).**

**Supplementary Fig. 3 QQ plot of MWAS using baseline MDS-UPDRS total score as a clinical trait.** As shown in the plot, almost all the CpGs on the diagonal of the plot and seldom the *P-*value of CpGs were smaller than expected, which indicates that little CpG was found to be significantly associated with this clinical trait.

**Supplementary Fig. 4 Heatmap of cell types associated with PD aggravation**. Abbreviations: Th1 cells, Type 1 T-helper cells; aDC, Activated dendritic cells; CD4+Tem, CD4+ effector memory T-cells; CD8+ Tcm, CD8+ central memory T cells; MSC, Mesenchymal stem cells; Th2 cells, Type 2 T-helper cells; HSC, Hematopoietic stem cells.

**Supplementary Fig. 5 Identifying drugs and potential drug targets for PD. a** L1000CDS^2^ analysis for drug discovery. DEGs in the pathway of regulating pluripotency of stem cells (KEGG ID: hsa04550) were submitted to L1000CDS2 to predict potential agents. DEGs overlapping with background genes of withaferin A in L1000 signatures were considered WFA-regulated genes. Five WFA-regulated genes were confirmed by the validation datasets. **b** Downregulated aggravator genes and PD drug targets in the neuron presynaptic terminal of dopaminergic synapse. The green color indicates drug targets of PD, while the magenta color indicates aggravator gene.

**Supplementary Fig. 6 Morphology and immunofluorescent identification of NSCs. a** Bright field image of suspended neurospheres at P1 (passage 1) that was cultured in complete StemPro^®^ NSC SFM for 6 days (Scale bar = 100 μm). **b** Bright field image of adherent NSCs at P2 (passage 2) that were cultured in complete StemPro^®^ NSC SFM (scale bar = 100 μm). **c** Immunofluorescent identification of NSCs with primary mouse anti-Nestin monoclonal antibody (ab6142, 1:100, Abcam, Cambridge, UK) (Scale bar = 100 μm). **d** Merged fluorescence image of nestin and DAPI (scale bar = 100 μm).

## Notes

### Competing Interest Statement

The authors have declared no competing interest.

