## Supplementary figures and images for "Interpreting the dynamic pathogenesis of Parkinson’s disease by longitudinal blood transcriptome analysis"

### Supplementary Figure 1

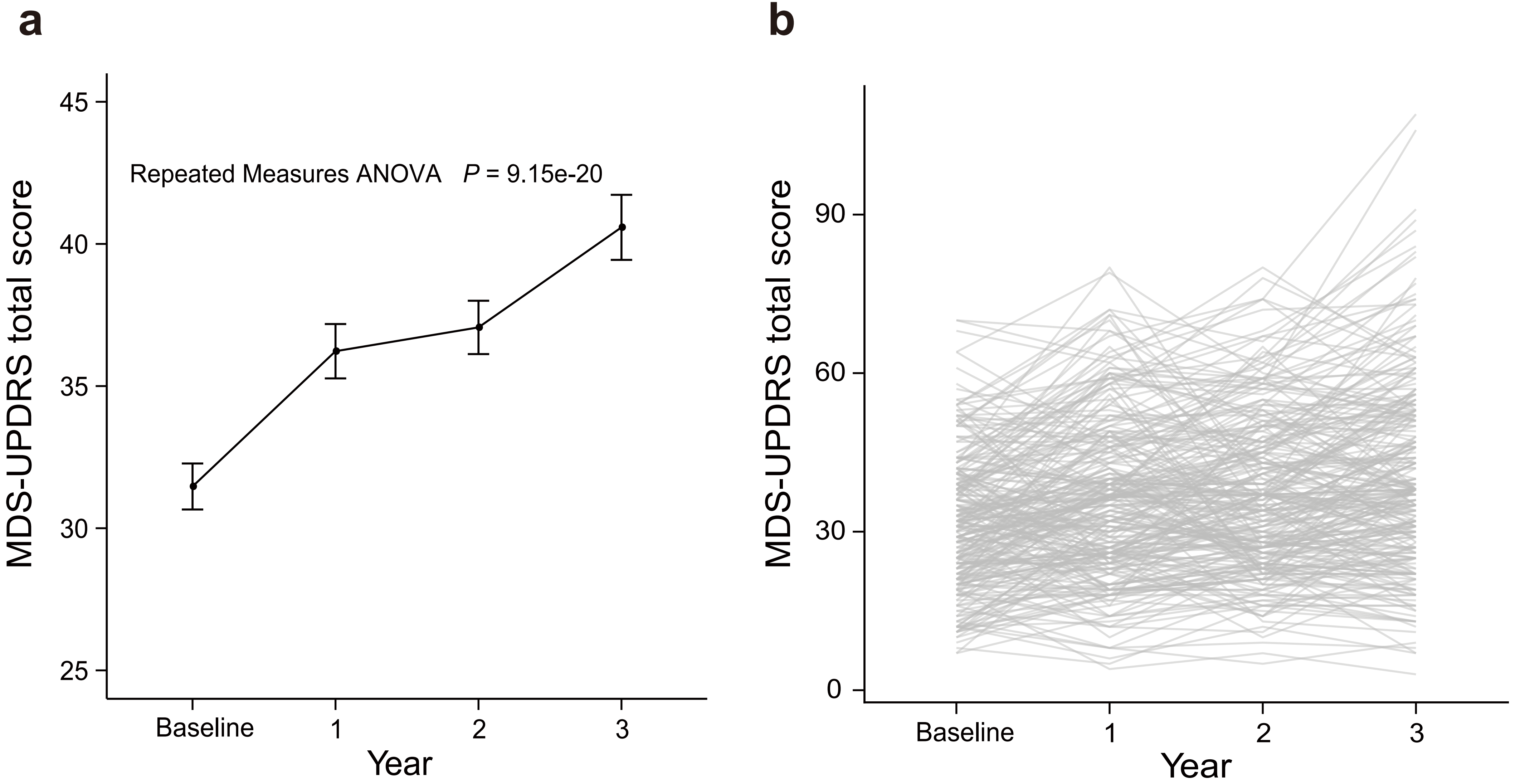

### Supplementary Figure 2

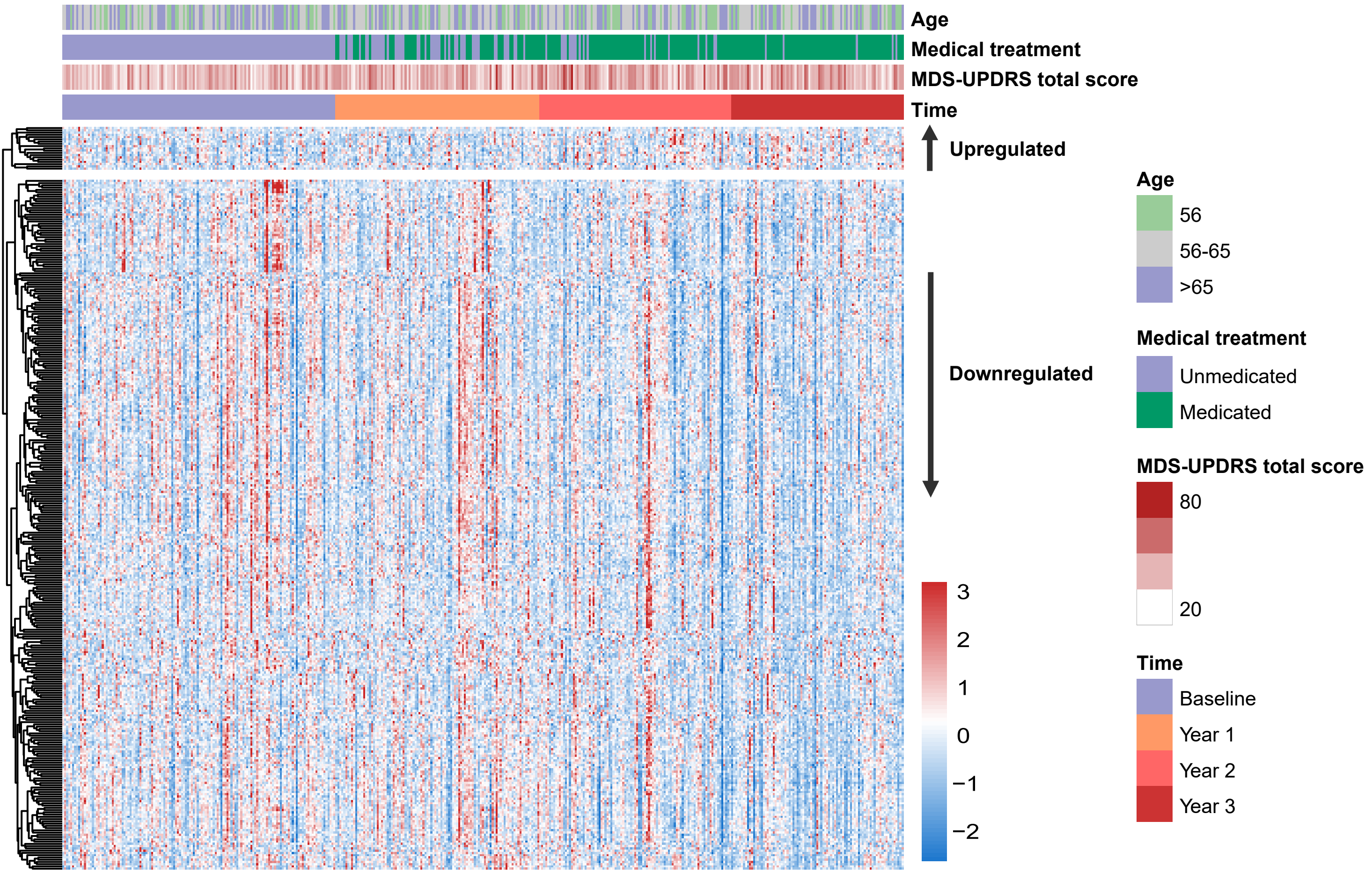

### Supplementary Figure 3

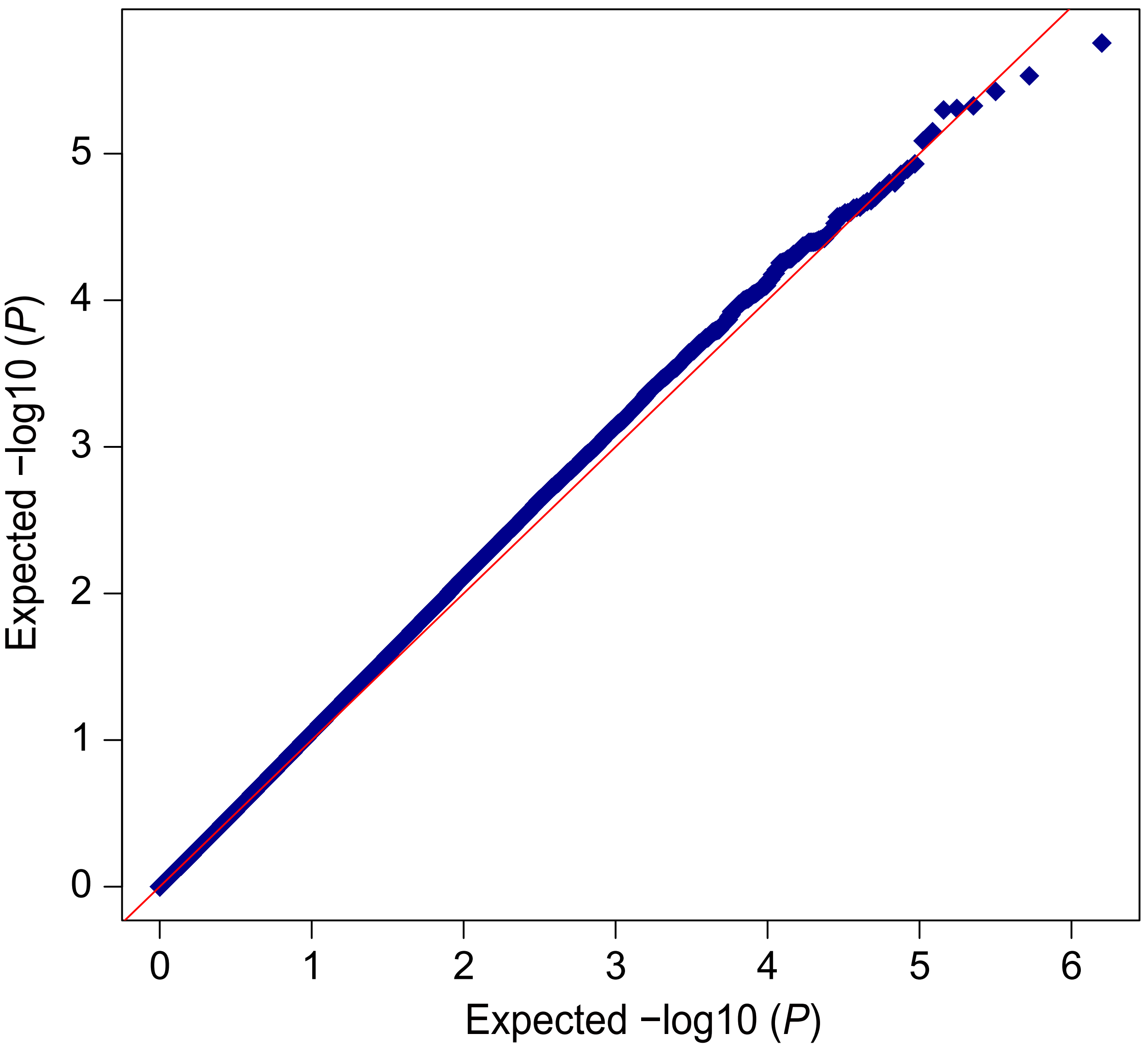

### Supplementary Figure 4

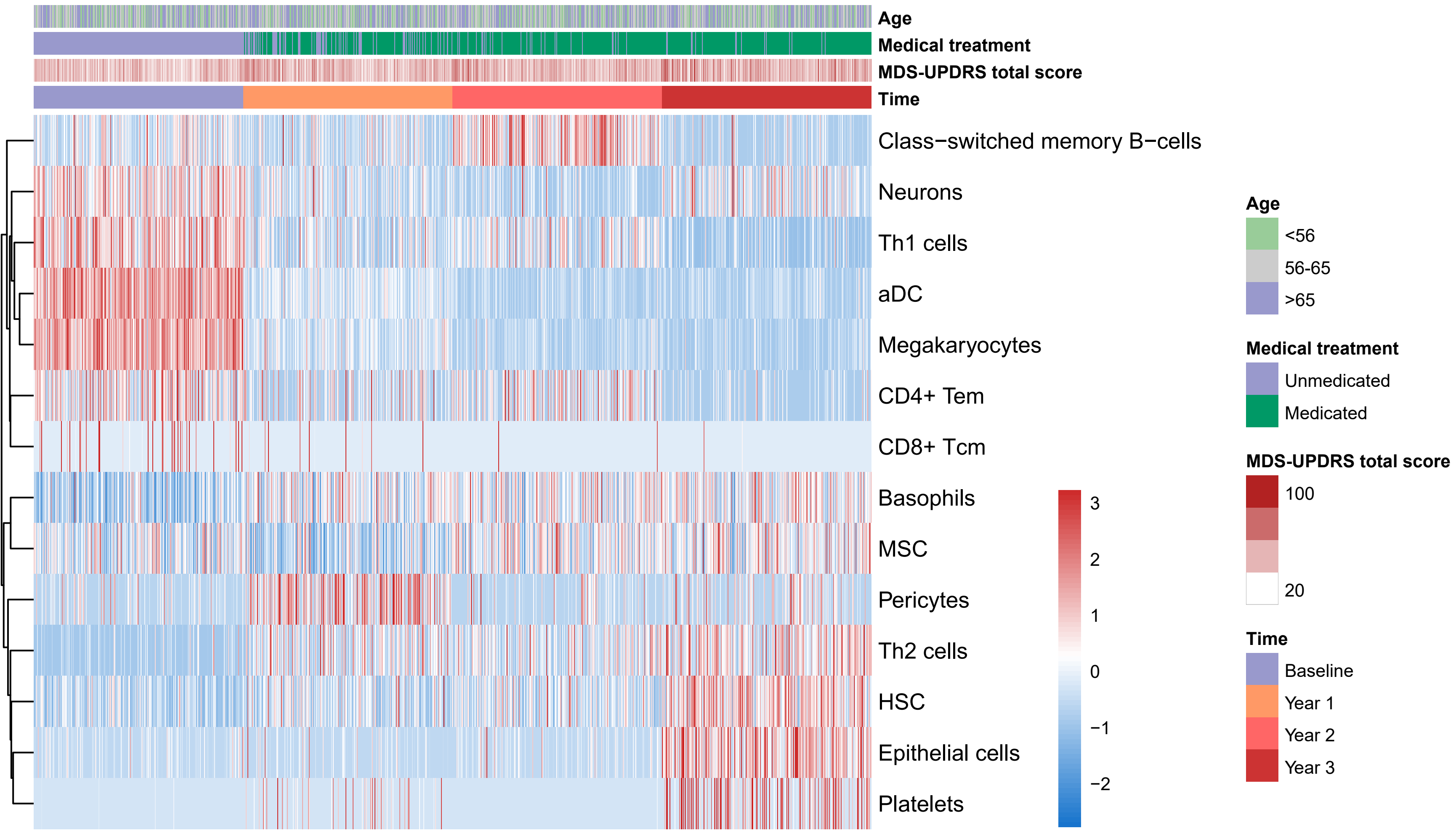

### Supplementary Figure 5

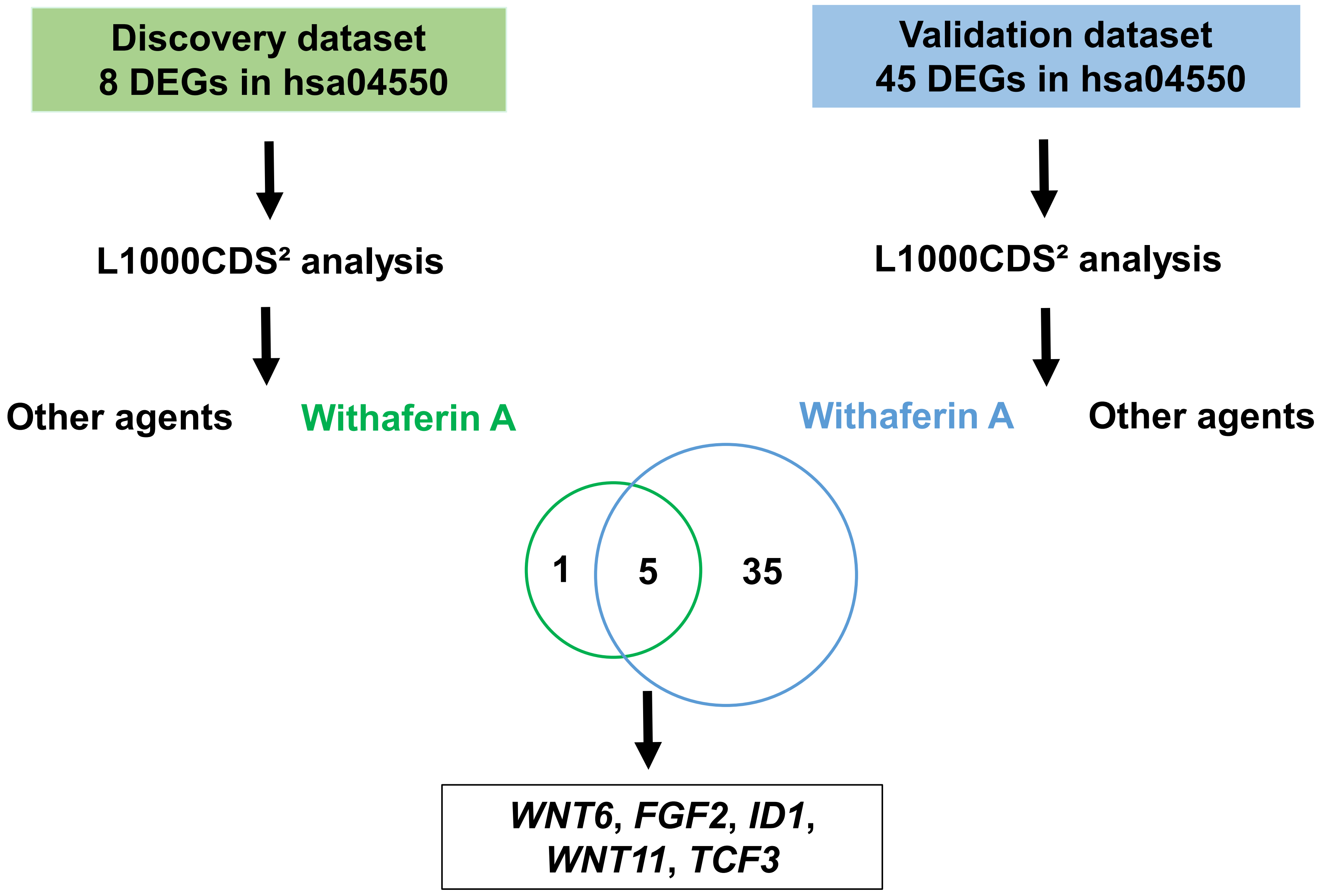

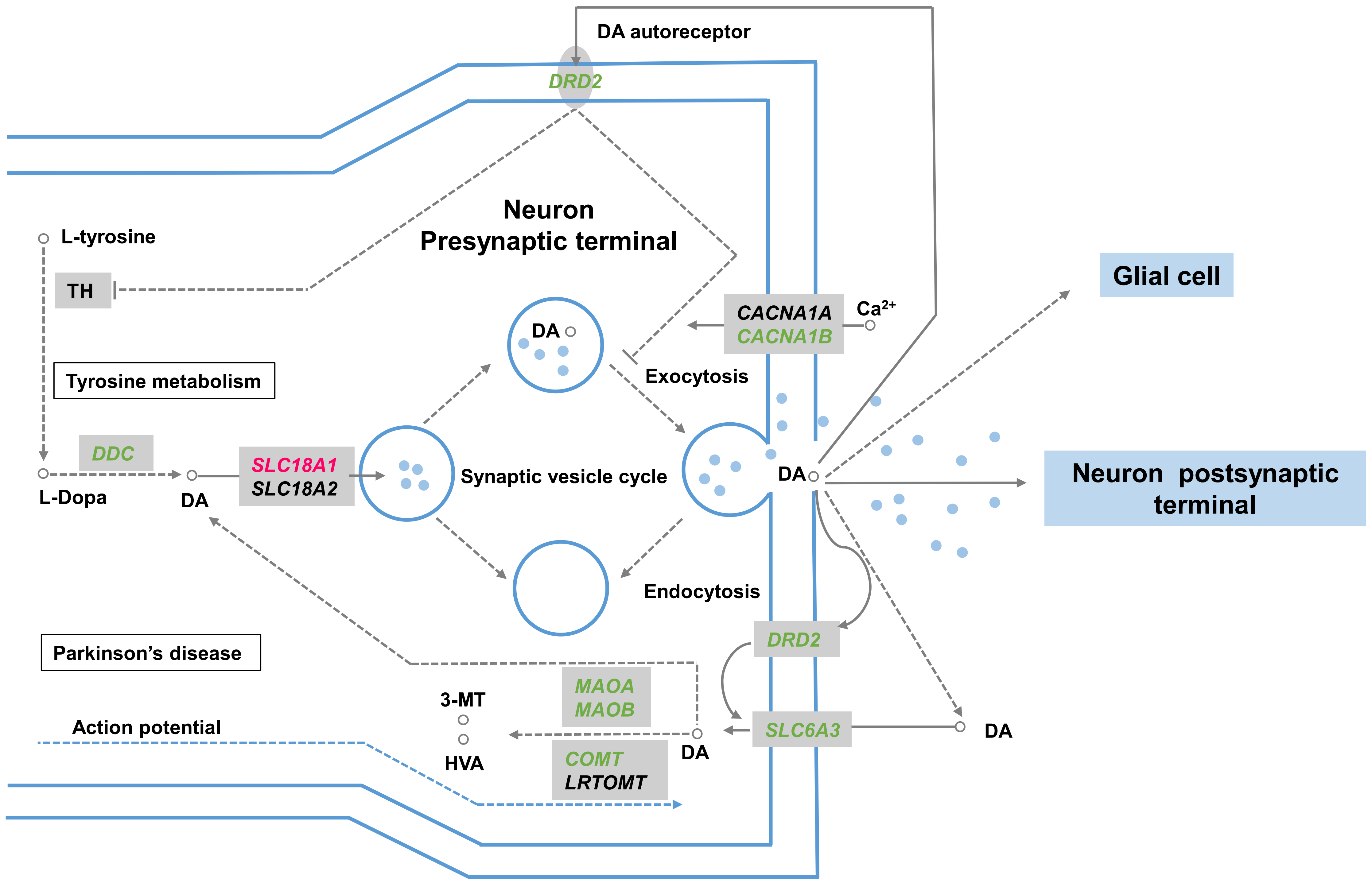

### Supplementary Figure 6

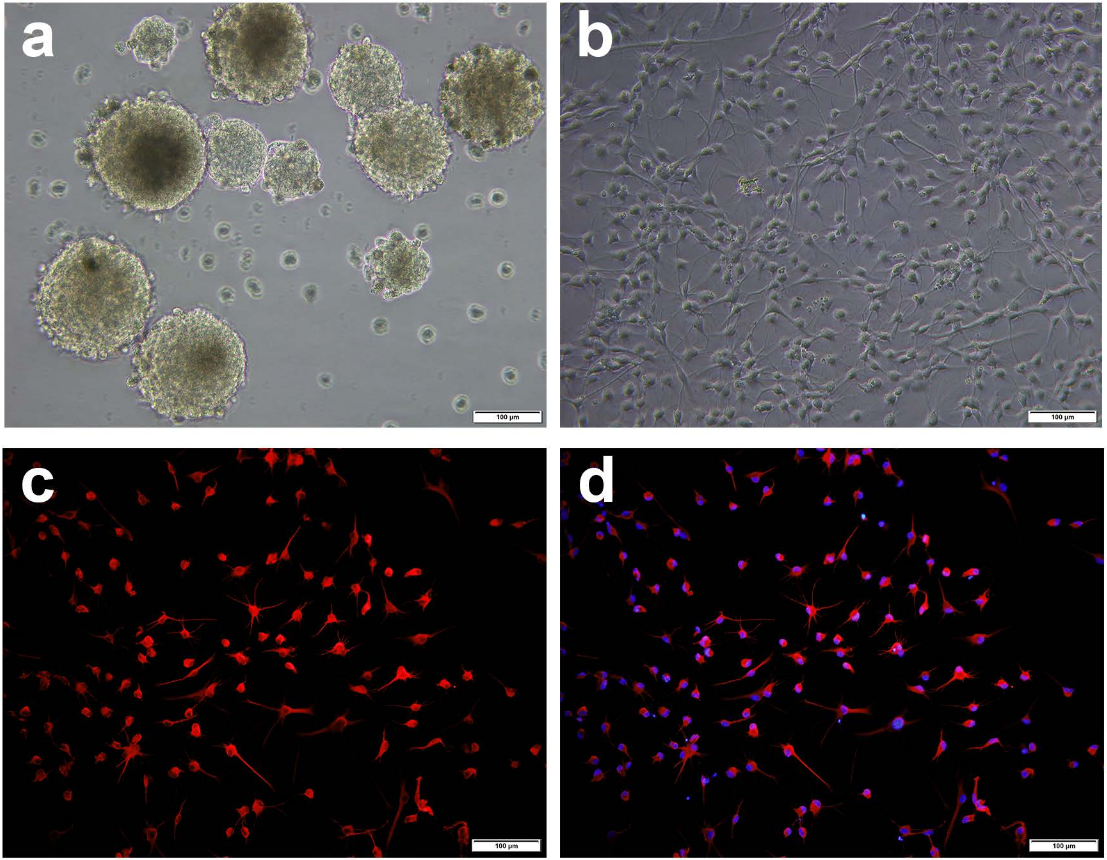
